# Inflammation is the main risk factor of voriconazole overdose in hematological patients

**DOI:** 10.1101/326595

**Authors:** Elodie Gautier-Veyret, A Truffot, S Bailly, X Fonrose, A Thiebaut-Bertrand, J Tonini, JY Cahn, F Stanke-Labesque

## Abstract

Aim: Voriconazole (VRC) overdoses are frequent and expose patients at high risk of adverse effects. This case-control study performed in hematological patients who benefited from VRC therapeutic drug monitoring from January 2012 to December 2015 aimed to identify risk factors of VRC overdose.

Methods: Pharmacogenetic, biological, and demographic parameters at the time of VRC trough concentration (Cmin) were retrospectively collected from medical records. Cases (VRC overdose: defined by a VRC Cmin ≥ 4 mg/l; n = 31) were compared to controls (no VRC overdose: defined by VRC Cmin < 4 mg/L; n = 31) using non-parametric or Chi-square tests followed by multivariable analysis.

Results: VRC overdoses were significantly associated with high CRP and bilirubin levels, intra-venous administration, and age in univariable analysis. In contrast, the proportion of CYP genotypes (CYP2C19, CYP3A4, or CYP3A5, considered alone or combined in a genetic score^1^) were not significantly different between patients who experienced a VRC overdose and those who did not. In multivariable analysis, the class of CRP level (defined by median CRP levels of 96 mg/l) was the sole independent risk factor of VRC overdose (p < 0.01). Patients with CRP levels > 96 mg/l had a 27-fold (IC 95%: [6-106]) higher risk of VRC overdose than patients with CRP levels ≤ 96 mg/l.

Conclusion: This study demonstrates that inflammatory status, assessed by CRP levels, is the main risk factor of VRC overdose in French hematological patients, whereas pharmacogenetic determinants do not appear to be involved.

## Introduction

Invasive fungal infections (IFI) are a major concern in immunocompromised patients. Among them, invasive aspergillosis (IA) is particularly feared in hematological units, as mortality occurs in 19 to 61% of cases (1). The management of antifungal therapy of IA is challenging due to various issues inherent either to inadequate exposure or drug safety. The first-line treatment of IA, voriconazole (VRC), is notably concerned by such issues, as this drug exhibits both highly variable pharmacokinetics and a narrow therapeutic range (2). Thus, VRC therapeutic drug monitoring (TDM) is recommended to optimize treatment (2, 3), as VRC trough concentrations (Cmin) are associated with both efficacy and toxicity (4).

Despite TDM, VRC Cmin are frequently out-of-the therapeutic range in clinical practice. Indeed, subtherapeutic VRC Cmin have been found in 20 to 33% of patients (5, 6) and supratherapeutic VRC Cmin in 8.8 to 23% of cases (5–7), depending on the therapeutic threshold and study population. Although subtherapeutic VRC Cmin are often explained by gain-of-function single nucleotide polymorphism (SNP) *17 for CYP2C19 (at least in the Caucasian population) (8–11), comedication by enzymatic inducers (12), or non-observance (9), the determinants of supratherapeutic VRC Cmin have been less investigated. Numerous factors, such as liver dysfunction, the use of enzymatic inhibitors (13), the presence of loss-of-function SNPs for CYP2C19 (14), and inflammation (15, 16), have already been identified to be associated with increased VRC Cmin, but their impact on the occurrence of VRC overdose has never been investigated. This issue is all the more relevant as 1) the magnitude of the effect of each individual factor on VRC Cmin appears to be relatively weak (4) and 2) some of these factors, such as inflammation and genetic variants of CYP2C19, may be linked (17–19) and interact (15, 16). Here, we aimed to identify risk factors associated with VRC overdose in a cohort of hematological patients treated by VRC followed by TDM.

## Materials and methods

### Study design

This retrospective case-control study was conducted at Grenoble Alps University Hospital, France in a hematological unit. Adult patients (> 18 years old) who experienced a VRC overdose, defined by VRC Cmin determined at steady-state ≥ 4 mg/l (2) between January 2012 and December 2015 were eligible. Patients were excluded in cases of prescription error, missing data, sampling before pharmacokinetic steady state, suspicion of non-observance, unavailability of their DNA and also if they were < 18 years (see Figure 1 for details).

**Figure 1.**
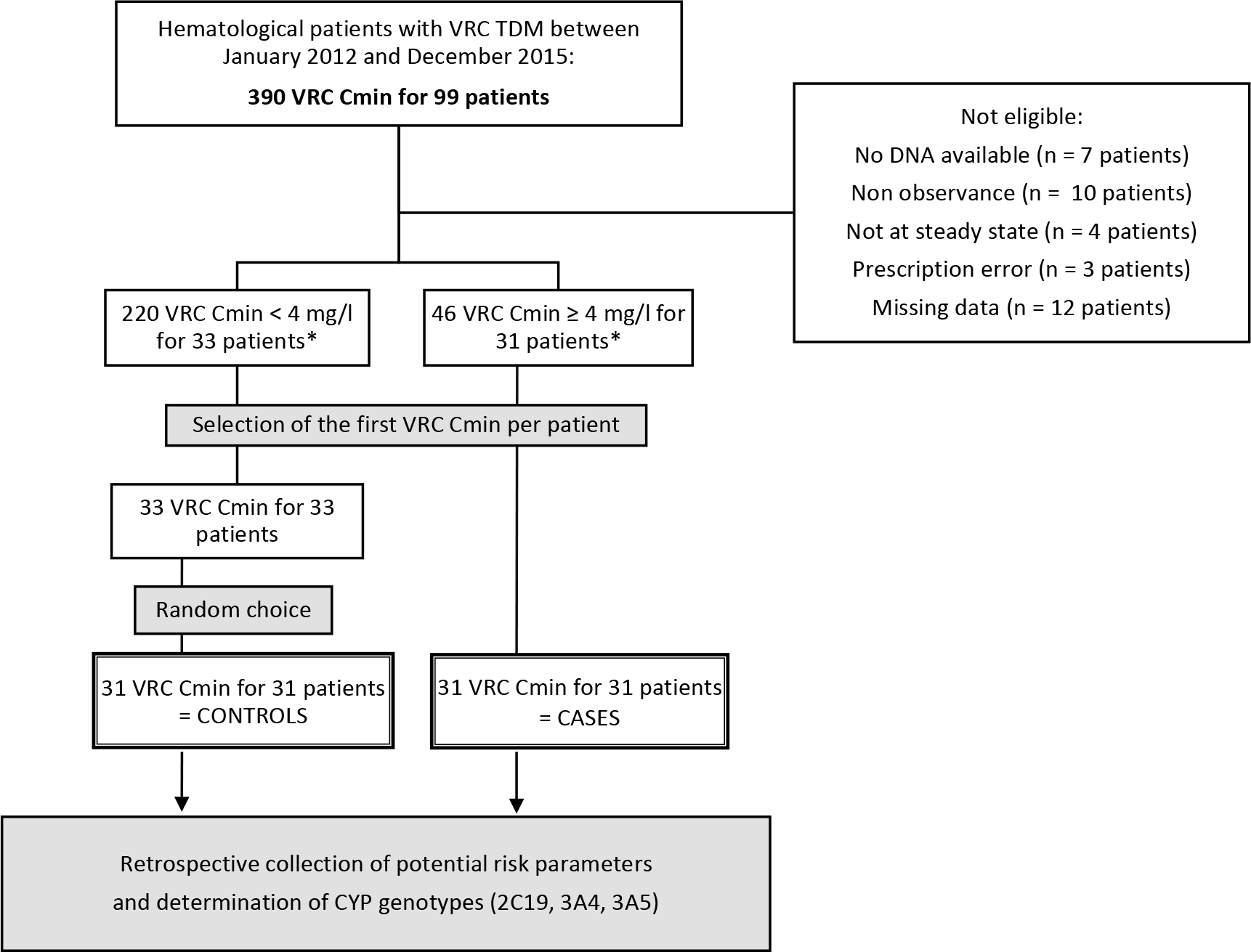
Flow chart. VRC, voriconazole; TDM, therapeutic drug monitoring; Cmin, trough concentration. * One patient with two treatment sequences was included in both groups.

Risk factors of VRC overdose were assessed by comparing patients experiencing VRC overdose to control patients (those with therapeutic VRC Cmin between 1 and 4 mg/l), who were randomly included among all patients being followed by VRC TDM during the same period. The inclusion and non-inclusion criteria for the selection of controls were similar to those for patients who experienced a VRC overdose, except for the VRC Cmin.

Demographic, clinical (underlying disease, hematopoietic stem cell transplantation), biological (CRP, ALAT, Bilirubin, VRC Cmin), and pharmaceutical data (VRC daily dose, route of administration, comedication with pump proton inhibitors (PPIs)) were retrospectively collected from medical records. If several VRC overdose episodes occurred for one patient, only the first was considered. The inflammatory status was assessed by determining the CRP level the same day as determination of the VRC Cmin. The absence of a strong enzymatic inducer or inhibitor was verified.

### Ethics statement

Informed written consent was obtained for all patients for genetic analysis, sample collection, and use of their data, in accordance with the Declaration of Helsinki. The study and biobank were approved by the IRB 6705 (CPP Sud Est 5, Grenoble, France).

### Therapeutic drug monitoring

Plasma VRC Cmin were determined at steady state for samples handled just before VRC administration using a validated liquid chromatography-tandem mass spectrometry (20).

### Genotyping and determination of the combined genetic score

Genotyping of CYP2C19 and CYP3A was performed on residual samples taken before allograft and stored in a biological sample collection (CRD-2013-1983). Only residual samples taken before allograft were considered for hematopoietic stem cell allograft recipients. The presence of CYP2C19*2 (rs4244285), CYP2C19*17 (rs72558186), CYP3A5*3 (rs776746), and CYP3A4*22 (rs35599367) was investigated using the TaqMan allelic discrimination assay (Life Technologies, Illkirch, France). The phenotype of CYP2C19 was defined according to the Clinical Pharmacogenetics Implementation Consortium (CPIC) guidelines for CYP2C19 and voriconazole (21). In addition, a genetic score, integrating both CYP2C19 and CYP3A genotypes, was determined for each patient according to our previous studies (11, 22): the higher the genetic score, the faster the metabolism of the patient. The Hardy-Weinberg equilibrium was tested for each polymorphism by the online method of Rodriguez *et al.* (23).

### Statistical analysis

Quantitative data are expressed as medians (25^th^-75^th^ percentiles). The link between VRC overdose and potential risk factors was investigated either by comparing the distribution of qualitative data (sex, cotreatment by PPI, route of administration, class of CRP based on a median CRP level of 96 mg/L, phenotype of CYP2C19, etc.) with either a Chi-square or Fisher’s exact test by comparing the values of continuous data (VRC dosing, CRP levels) by the Man-Whitney test. A multivariable logistic regression model was performed to account for potential confounders. CRP was categorized into two classes due to the absence of log-linearity: CRP ≤ or > 96 mg/l, based on the median. Variables with a p value threshold of 0.2 in univariable analysis were selected and introduced in the multivariable analysis, except in cases of collinearity. A stepwise procedure was used to select the final model. Statistical analyses were performed using SAS v9.4 (SAS Institute Inc., Cary, NC). Significance was considered for p < 0.05.

## Results

Baseline characteristics of patients and potential risk factors are described in Table 1. Thirty-one patients, who experienced a VRC overdose, as well as thirty-one controls without a toxic VRC Cmin, were included. The median for the VRC Cmin in the cases (patients who experienced a VRC overdose) was 5.3 mg/l *versus* 1.0 mg/l for the control group (Table 1).

**Table 1.**
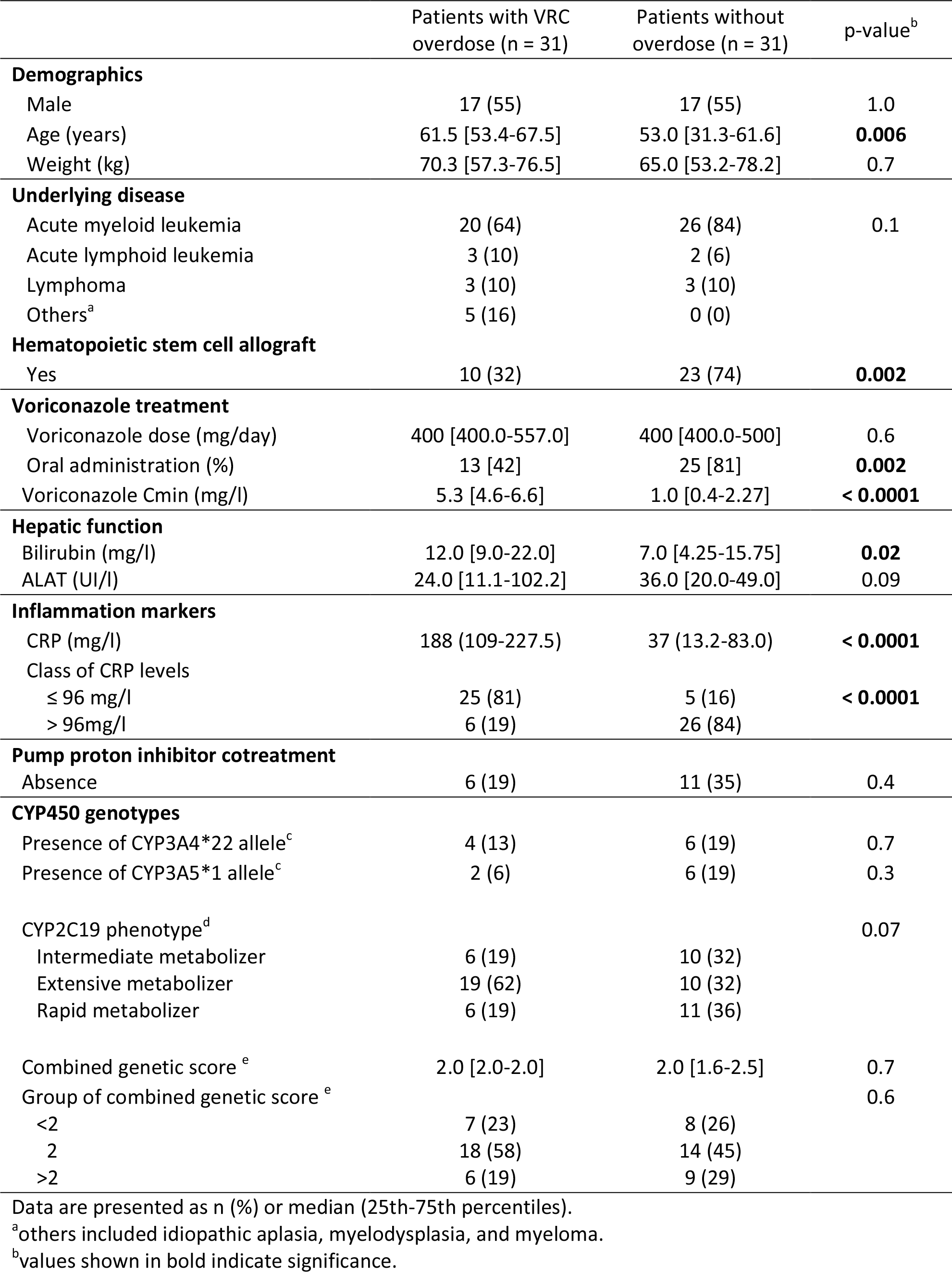
Link between VRC overdose and potential risk factors.

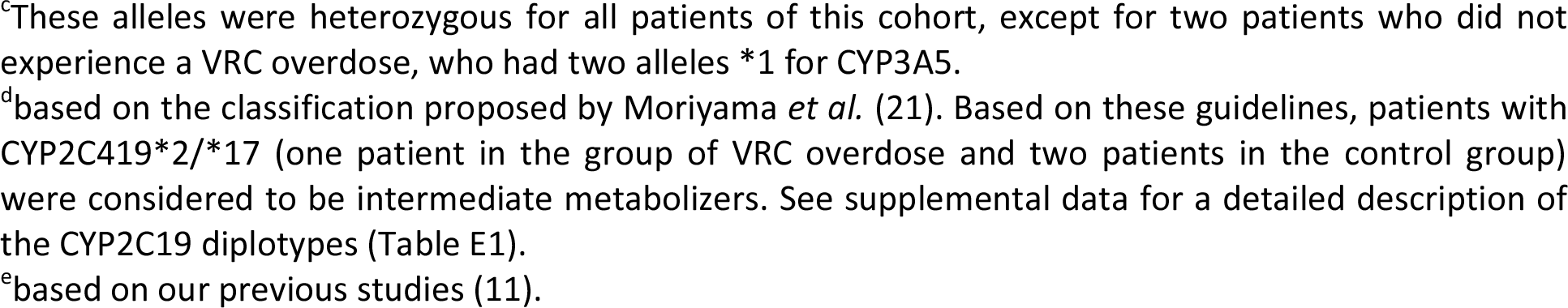

In univariable analysis, patients in the case group were older, exhibited higher bilirubin levels, and were more frequently treated by intravenous voriconazole than those in the control group (Table 1). The proportion of allografted patients was lower in the case than control group. In addition, the distribution of CRP levels was significantly different between patients who experienced a VRC overdose and those who did not (table 1), with higher CRP levels for cases than controls (Figure 2).

**Figure 2.**
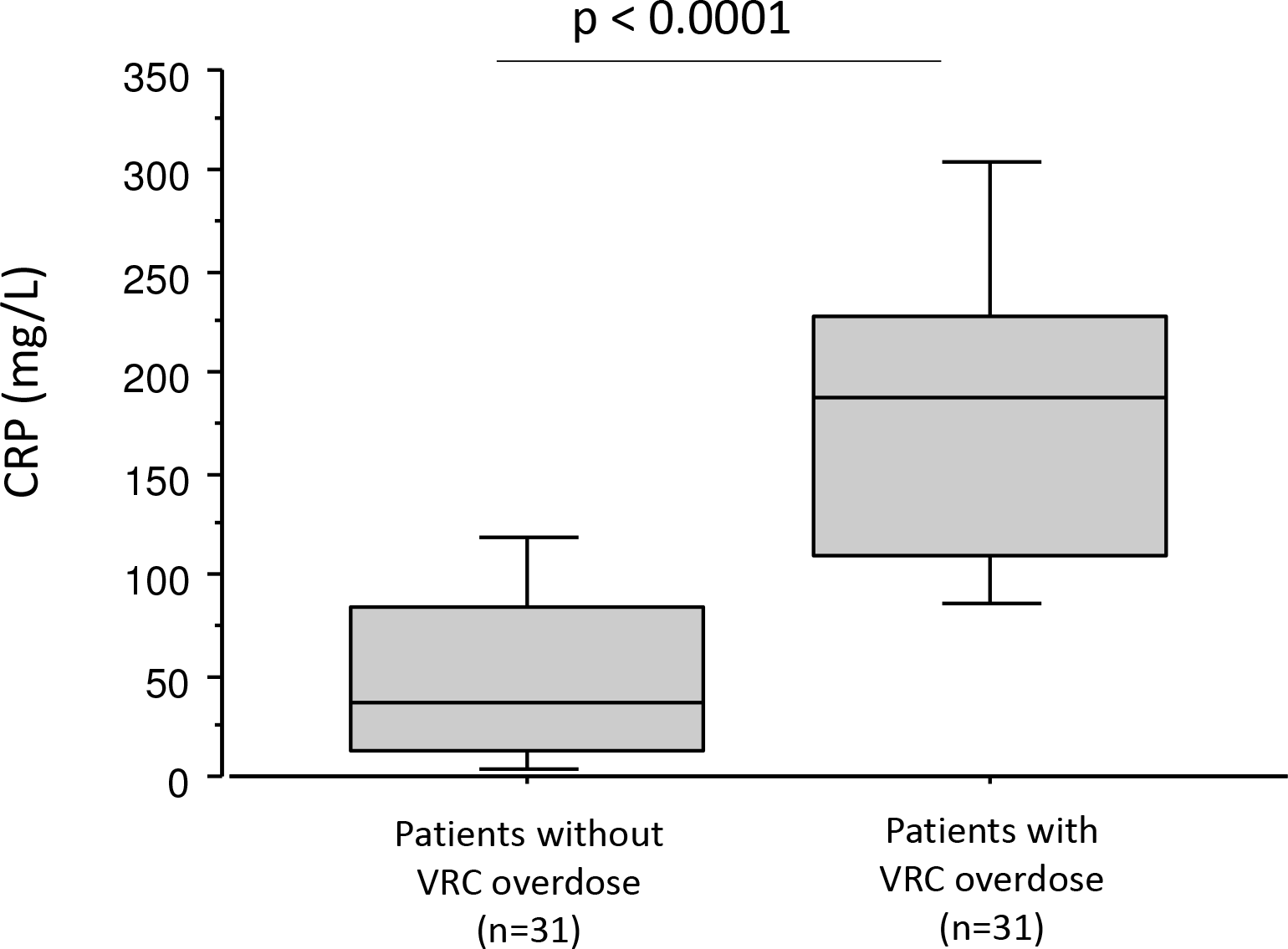
C reactive protein levels in patients who experienced a voriconazole overdose and those who did not.

In contrast, VRC dose, sex, weight, PPI treatment, ALAT levels, underlying disease, and pharmacogenetic parameters were not associated with a risk of VRC overdose, except for the distribution of the CYP2C19 phenotype (table 1) or diplotype (Table E1), which tended to differ between patients who experienced a VRC overdose and those that did not (p = 0.07 and p = 0.1, respectively). There was no association between pharmacogenetic parameters and risk of VRC overdose, either when cytochromes CYP2C19, CYP3A4, or CYP3A5 were considered alone or evaluated in the combined genetic score (table 1).

We investigated a possible link between CRP levels and pharmacogenetic parameters (17–19) by comparing CRP levels (considered as continuous variables or CRP classes) according to CYP2C19 phenotype and combined genetic score class. There was no difference in CRP levels depending on either CYP2C19 phenotype or combined genetic score class (Figure E1 and Table E2).

In the multivariable analysis, only the class of CRP level was significantly and independently associated with VRC overdose (p < 0.01). Patients with a CRP > 96 mg/l had a greater risk of VRC overdose than patients with a CRP ≤ 96 mg/l (odds ratio (95 % confidence interval): 27 [6-106]).

## Discussion

This study identified, for the first-time, inflammation (assessed by CRP levels) to be a strong independent risk factor of VRC overdose in hematological patients. These data suggest that hematological patients may be at increased risk of supratherapeutic VRC Cmin and associated toxicity when they exhibit increased CRP levels, which is very frequent for patients with IA (24, 25).

Such an elevated CRP-level associated risk of VRC overdose is in accordance with several previous studies reporting positive associations between elevated CRP levels and high VRC Cmin (15, 16, 26–28). This is probably related to inflammation-induced phenoconversion, during which proinflammatory cytokines, such as interleukin-6, reduce both CYP3A4 and CYP2C19 expression and activity (29). Hence, it is probable that inflammation rapidly inhibits VRC metabolism by reducing CYP3A4 and CYP2C19 activities, which ultimately results in increased VRC Cmin and greater risk of overdose.

PPI cotreatment, advanced age, and high bilirubin levels, which have already been reported to be associated with increased VRC Cmin, were not associated (PPI treatment) or only associated in univariable analysis (age and hepatic dysfunction) with a risk of VRC overdose in this study. Similarly, none of the pharmacogenetic parameters, including the CYP2C19 phenotype or the CYP3A4 and CYP3A5 genotypes, considered alone or in a combined genetic score, were associated with VRC overdose. This result is surprising, as numerous previous studies have demonstrated that the loss-of-function SNPs affecting CYP2C19 (*2 or *3) (30–32) or CYP3A4 (*22) (11, 33–36) were associated with elevated VRC Cmin. These discrepancies may be explained by the relatively low number of patients included, which limited the statistical power of our study and increased the width of the confidence intervals. Despite this limitation, our work clearly demonstrates that inflammation, shown by high CRP levels, is the sole independent risk factor of VRC overdose in French hematological patients, completely masking the effects of other potential risk factors.

In addition, we investigated the possible link between CRP levels and the CYP2C19 phenotype by additional statistical analyses (see Table E2 and Figure E1 in supplemental data). Although there was no association between CRP and pharmacogenetic parameters (both CYP2C19 phenotype and class of combined genetic score), the absence of a link cannot be completely ruled out as CRP and IL-6 levels have been reported to be elevated in poor CYP2C19 metabolizers (17–19). These findings suggest that an association between loss-of-function SNPs of CYP2C19 and elevated VRC Cmin (8, 30, 37) could result in reduced VRC metabolism due to diminished CYP2C19 activity (direct effect), as well as an increase in IL-6 levels, and hence down-regulation of CYP2C19 and CYP3A4 activities (indirect effect). Further studies should be performed to clarify the respective roles of inflammation and pharmacogenetic determinants on VRC Cmin and explore their eventual link, especially as the effect of inflammation on VRC Cmin may be modulated by pharmacogenomic parameters (15, 38).

This study had several limitations. First, it was retrospective, performed in a single center, and only a small number of subjects was included. Second, data on the clinical consequences of VRC overdose in terms of side effects, treatment discontinuation, and efficacy of treatment was absent. This aspect may be particularly important to explore in the future, as studies have already shown that elevated pro-inflammatory cytokine levels are a predictor of poor outcome in invasive aspergillosis (24, 25).

In conclusion, our study identified inflammation as the main risk factor of VRC overdose in French hematological patients, whereas pharmacogenetic determinants do not appear to be involved. Hence, the inflammatory status, evaluated by CRP levels, must to assessed for every VRC overdose and individualized dose adjustments will probably need to be integrated during the longitudinal evolution of inflammation (26, 39) to obtain a VRC Cmin within the therapeutic range. In the era of individualized VRC therapy based on CYP2C19 phenotype (21), a more global approach may be of interest to effectively and rapidly individualize VRC therapy, taking into account not only the CYP2C19 phenotype (21), but also other determinants of VRC Cmin, particularly SNPs affecting other enzymes involved in VRC metabolism, such as CYP3A4 (11, 33–36), eventual comedication by enzymatic inhibitors/inducers (11, 40) and, of course, inflammatory status (15, 16, 26–28).

## Acknowledgments

The authors thank Christel Roche and Mélanie Davallet-Pin, Cécile Pistone-Girard and Karine Scalabrino for their excellent technical assistance.

**Conflict of interest:** none

## Additional data

**Table E1.**
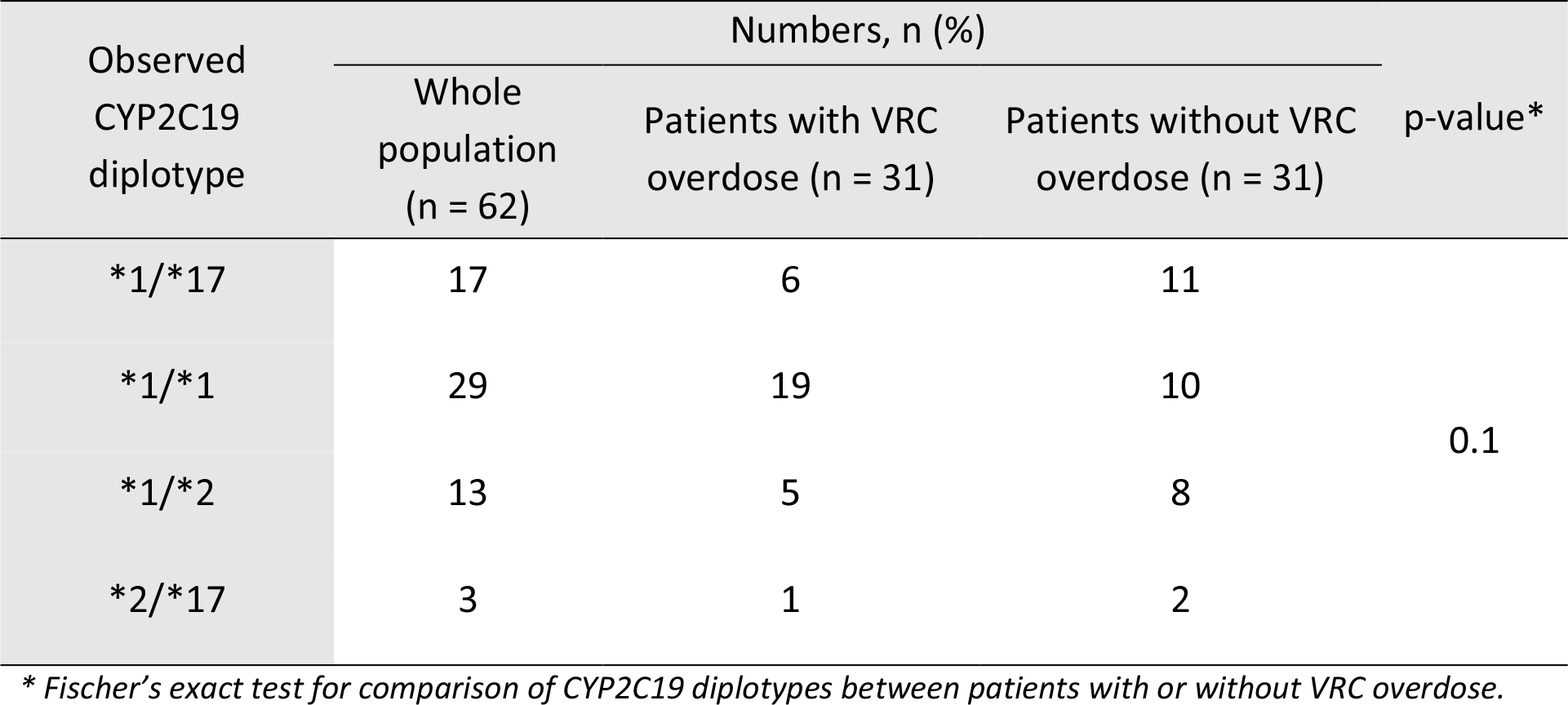
Distribution of CYP2C19 genotypes score

| Observed<br>CYP2C19<br>diplotype | Numbers, n (%) |  |  | p-value* |
| --- | --- | --- | --- | --- |
|  | Whole<br>population<br>(n = 62) | Patients with VRC<br>overdose (n = 31) | Patients without VRC<br>overdose (n = 31) |  |
| *1/*17 | 17 | 6 | 11 | 0.1 |
| *1/*1 | 29 | 19 | 10 |  |
| *1/*2 | 13 | 5 | 8 |  |
| *2/*17 | 3 | 1 | 2 |  |

\* Fischer's exact test for comparison of CYP2C19 diplotypes between patients with or without VRC overdose.

**Table E2.** Distribution of CRP levels according to CYP2C19 phenotype and class of combined genetic score

|  | Class of CRP levels |  | P value* |
| --- | --- | --- | --- |
|  | Class 1<br>(≤ 96 mg/L) | Class 2<br>(> 96 mg/l) |  |
| **CYP2C19 phenotype**<sup>1</sup> |  |  |  |
| Intermediate metabolizer | 5 | 8 | 0.3 |
| Extensive metabolizer | 14 | 15 |  |
| Rapid metabolizer | 11 | 6 |  |
| **Class of combined genetic score**<sup>2</sup> |  |  |  |
| < 2 | 6 | 10 | 0.5 |
| 2 | 15 | 16 |  |
| > 2 | 9 | 6 |  |

CRP, C reactive protein. \*Chi<sup>2</sup> exact test for comparison of classes of CRP levels according to CYP2C19 phenotype and class of combined genetic score

---

**Figure E1.**
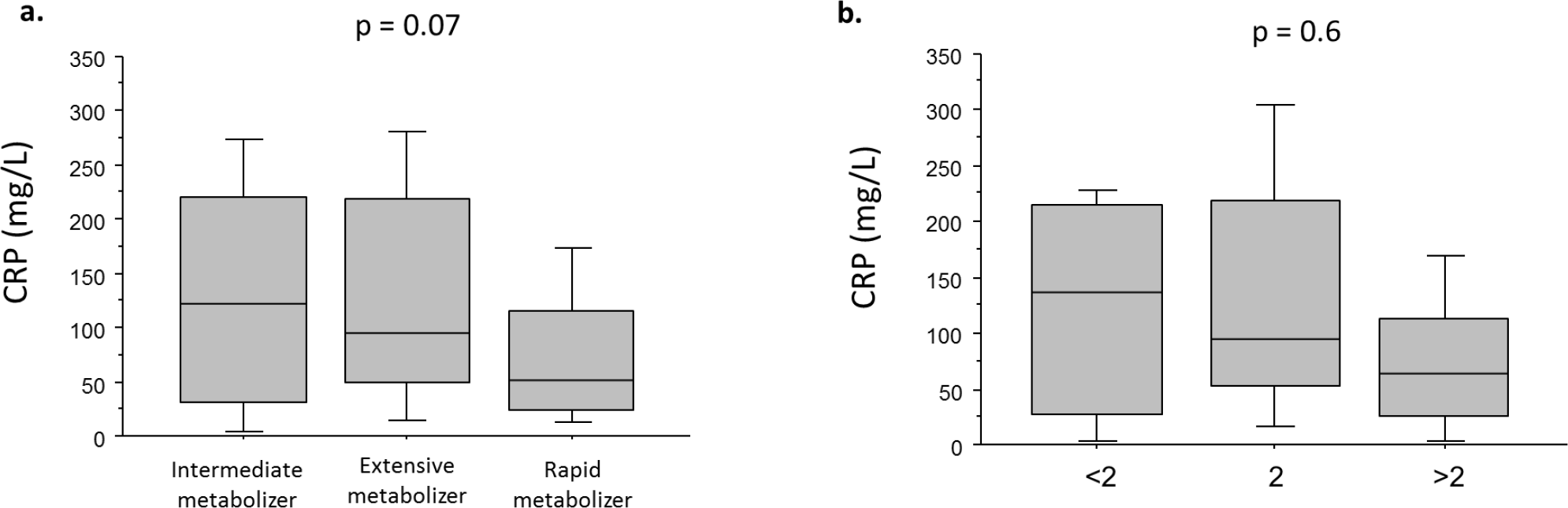
C reactive protein levels according to CYP2C19 phenotype (a) and class of combined genetic score (b)

## Footnotes

1 Gautier-Veyret et al., AAC 2015

